# Molecular Evolution under Macro-Perturbation Barrier and the Fixation Process of Polyploidization

**DOI:** 10.1101/2025.11.19.689354

**Authors:** Xun Gu

## Abstract

The Neutral Theory in molecular evolution mainly addresses the allele frequency change of individual loci, postulating that the fixation process of a mutant is independent of other loci. For instance, the basic formula of molecular evolution claims that the rate of molecular evolution (λ) is determined by the mutation rate (*v*), the coefficient of selection (*s*) and the effective population size (*N*_*e*_), asserting the basic rule of molecular evolution: λ>*v* if *s*>0 (positive selection), λ=*v* if *s*=0 (neutrality) or λ<*v* if *s*<0 (negative selection). However, many studies have indicated that this independent assumption should be examined carefully. This paper studies focused on the effect of major-perturbation barrier on molecular evolution, which refers to a rare, randomly occurred major genetic, epigenetic or environmental event that virtually stopped the fixation process of a mutation, resulting in loss of the mutation. A special diffusion model called KAC model was invoked, which allows the stochastic trajectory of gene frequency toward fixation can be randomly stopped at any time with a certain rate. The analytical form of the rate of molecular evolution showed that a strictly neutral mutant would evolve more slowly than the mutation rate due to the macro-perturbation barrier. A further quasi-neutrality analysis (the rate of molecular evolution equals to the mutation rate) was carried out, based upon the balance between the selection advantage of the mutant and the macro-perturbation barrier. Finally, the theory of macro-perturbation barrier was illustrated by the fixation process of polyploidization, as the outcome of the short-term selective advantages and the genome instability.

## Introduction

In the earliest formulations of the Neutral Theory, researchers focused on the allele frequency change of individual loci (genes or nucleotide sites), postulating that the process is independent between loci. This assumption is virtually equivalent to the principle of quasi-independence proposed by Lewontin (1974), that is, under selection, a character may evolve as if it is independent of other characters, although these characters are functionally related. Many theoretical results of population genetics are therefore mathematically tractable and empirically applicable. For instance, Kimura (1962) depicted how the rate of molecular evolution (λ) between species is determined by the mutation rate (*v*), the coefficient of selection (*s*) and the effective population size (*N*_*e*_), asserting the basic rule of molecular evolution: λ>*v* if *s*>0 (positive selection), λ=*v* if *s*=0 (neutrality) or λ<*v* if *s*<0 (negative selection).

For over decades, the basic formula of molecular evolution (Kimura 1962) has played a central role in the long-term neutralist-selectionist debate (Kimura 1968; 1983; Ohta 1973; Gillespie 1989; Hahn 2008; Ken and Hahn 2018; Jensen et al. 2019). It has been also recognized that some of arising controversies arising are partially due to the independent assumption between loci. Tentatively, interactions between loci can be classified into two categories. The first category is the genetic hitchhiking at linked loci. Regardless of the relative importance, two forms of hitchhiking, selective sweeps or background selection, tend to reduce the genetic diversity in linked nucleotide sites. Yet, it is important to note that neither of them affects the probability of fixation of neutral mutations (Birky and Walsh 198; Gillespie 2001). Hence, it is practically legitimate to study the rate of molecular evolution without considering the hitchhiking. The second category is the epistasis or the background effect. In spite of many underlying mechanisms, one may collectively called the perturbation effect in molecular evolution. There are two types of perturbations. The micro-perturbation results in slight fluctuation of selection among individuals who carry the mutation. Recently, Gu (2025a, 2025b) studied this problem extensively under the FSI (fluctuating selection among individuals) framework. Meanwhile, the macro-perturbation is caused by a rare, randomly occurred major genetic, epigenetic or environmental event that virtually stopped the fixation process of a mutation. The macro-perturbation barrier refers to the case when the ultimate fate of the mutation is being lost, whereas the macro-perturbation tunnel refers to the case when the ultimate fate of the mutation is being fixed.

Many factors, adaptive or non-adaptive, may contribute to the impact of macro-perturbation on the pace of molecular evolution. However, attempts to accommodate those underlying mechanisms in the derivation of the rate of molecular evolution have been shown difficult, and therefore, an effect-model approach would be desirable. This article addresses the problem of macro-perturbation barrier that decreases the rate of molecular evolution: invoking a special diffusion model called KAC model, which allows the stochastic trajectory of gene frequency toward fixation can be randomly stopped at any time with a certain rate. The analytical form for the rate of molecular evolution is derived, demonstrating that the rate of a strictly neutral mutant would be less than the mutation rate under the macro-perturbation barrier. We then analyze the quasi-neutrality, i.e., the rate of molecular evolution equals to the mutation rate, based upon the premise that the arising quasi-neutrality is due to the balance between the selection advantage of the mutant and the macro-perturbation barrier. Finally, the theoretical formulation of macro-perturbation barrier is applied to illustrate the real evolutionary scenario of the fixation process of polyploidization: a balance between the short-term selective advantages and the genome instability.

## Results and Discussion

### Molecular evolution as fixation process

Let *λ* be the rate of nucleotide substitution. In population genetics, fixation models attempt to describe how a new mutation can spread in a population, ultimately toward being fixed rather than being lost, due to either selection or genetic drift (Kimura 1962; 1983); one may see McCandlish and Stoltzfus (2014) for a comprehensive review. Here mutations are usually described as random events that have a rate of occurrence (the mutation rate *v*) and a contribution to fitness (the selection coefficient *s*). Suppose we have a diploid population of size *N* (the census population size). Then the total amount of such mutations appear in the population as a whole is simply *2Nv*. Since only a fraction of these mutations will eventually go to fixation, to calculate the substitution rate (λ), we need to multiply *2Nv* with the fixation probability of a new mutation (Kimura 1962). Let *u*(*p*) be the fixation probability of a mutation in the population, with the initial frequency *p*. Then, the formula of substitution rate can be concisely written as follows

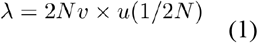

where the initial frequency set by 1/(2*N*) is based on the assumption of rare, single *de novo* mutation event. In addition to the initial frequency, the fixation probability is highly dependent of the selection coefficient (*s*) of the mutation: it becomes high when the mutation is beneficial (*s*>0) or low when the mutation is deleterious (*s*<0).

The key step to solve Eq.(1) is to derive the analytical form of fixation probability. Under the Wright-Fisher additive model, Kimura (1962) derived the well-known formula of molecular evolution, which depicts how the substitution rate (λ) between species is determined by the mutation rate (*v*), the coefficient of selection (*s*) and the effective population size (*N*_*e*_), that is,

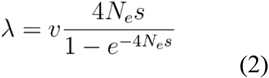

Eq.(2) asserts the basic rule of molecular evolution: λ>*v* if *s*>0 (positive selection), λ=*v* if *s*=0 (neutrality) or λ<*v* if *s*<0 (negative selection).

### Macro-perturbation barrier in the fixation process

Formally, a process is called a diffusion with a killing (also called KAC process) if the sample paths, denoted by *X*(*t*), behave like those of a regular diffusion until a possibly random time ζ when the process is killed. One may write a diffusion with a killing in the form {*X*(*t*), 0≤*t*<ζ}. It appears that a regular diffusion (without killing) is a special case of ζ=∞. For a diffusion with killing, at each point *x* there is a probability *k*(*x*)*dt*+*o*(*dt*) that the process ceases (is killed) over the infinitesimal time duration (*t, t+dt*), and probability 1-*k*(*x*)*dt*+*o*(*dt*) that no killing occurs. In symbols, we have

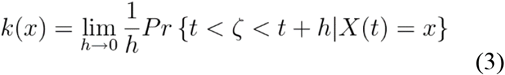

which is nonnegative, bounded and continuous. Intuitively, a killing function, *k*(*x*), is the probability of being killed at time *t* when the allele frequency is *x*.

We focus on the fixation process of a mutation in a given population. A phenomenon called macro-perturbation barrier refers to a rare, randomly occurred major genetic, epigenetic or environmental event that virtually stopped the fixation process of a mutation, which can be modeled by a fixation process with a killing. Specifically, we shall scrutinize the effect of macro-perturbation barrier by calculating to what extent the fixation probability of a mutant would be reduced by the associated killing process. Let *u*(*x*) be the probability of an allele to be ultimately fixed in the population, given the initial allele frequency (*x*). Under the KAC diffusion model, it is defined by

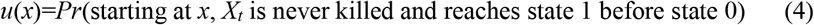

Following the derivation given by Karlin and Taylor (1981), one can show that the backward equation of *u*(*x*) with the KAC process is given by

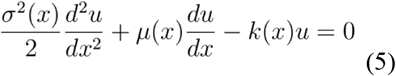

with the boundary conditions *u*(0)=0 and *u*(1)=1.

### Fixation probability under Wright-Fisher model with KAC

Consider two alleles at a locus, the wide-type (*A*) and the mutant (*a*) in a haploid population. The relative fitness of three genotypes *AA, Aa* and *aa* are additive, given by 1, 1+*s*, and 1+2*s*, respectively, where *s* is the coefficient of selection of mutant *a*. The allele frequency of the mutant *a* in a population changes across generations (*t*), jointly determined by the selection pressure (*s*), the random sampling of individuals in a finite population (the genetic drift), as well as the macro-perturbation. Under the Wright-Fisher additive model, the instantaneous mean *μ*(*x*) and the instantaneous variance *σ*^*2*^(*x*) are given by

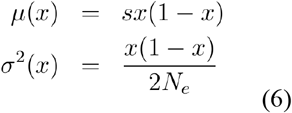

respectively, where *N*_*e*_ is the effective population size. Moreover, the killing function with respect to the fixation process can be written as follows

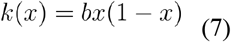

where *b* is the coefficient of macro-perturbation barrier. One may intuitively interpret Eq.(7) as follows. In addition to the coefficient *b* representing the arising rate of macro-perturbation barrier per generation, the probability that stops the fixation process is proportional to *x*, while the efficiency of KAC is proportional to 1-*x*.

Plugging Eq.(6) and Eq.(7) into Eq.(5), one can show that Eq.(5) can be simplified as follows

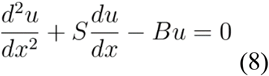

where S = 4*N*_*e*_*s* is the selection intensity, and *B* = 2*N*_*e*_*b* be the strength of macro-perturbation barrier. With the boundary conditions *u*(0)=0 and *u*(1)=1, we derive the analytical form of *u*(*x*) as follows

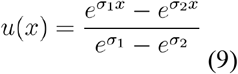

where two eigenvalues, *σ*_1_ and *σ*_2_, are given by

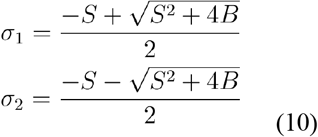

respectively. As *B*= 0, one can show *σ*_1_ = 0 and *σ*_2_ = −*S*, and so Eq.(9) is reduced by

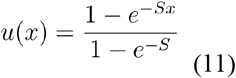

which is the well-known formula under the standard Wright-Fisher model.

### Rate of molecular evolution under macro-perturbation barrier

It appears that macro-perturbation barrier reduces the fixation probability and so as the rate of molecular evolution. When the initial frequency *x* is very small, the fixation probability given by Eq.(9) can be approximated by

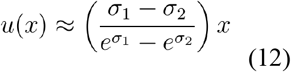

Therefore, the rate of molecular evolution defined by Eq.(1) can be written as follows

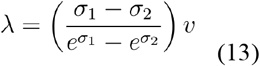

To facilitate further analysis, we define 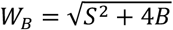 that measures the magnitude of joint effects of the selection and the macro-perturbation barrier. It follows that

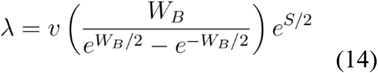

Eq.(14) shows that, in additional to the mutation rate (*v*), the rate of molecular evolution is determined by the selection intensity (*S*) and the magnitude (*W<SUB>B</SUB>*) of the joint effects. Fig.1 panel A presents the ratio of evolutionary rate to the mutation rate (*λ/v*) plotted against *B*, given the selection intensity *S*=-1, *S*=0, or *S*=1, respectively. Meanwhile, panel B presents the evolution-mutation rate ratio plotted against the election intensity (*S*), given the macro-perturbation barrier *B*=0, *B*=1 or *B*=5, respectively. Overall, macro-perturbation barrier is to decelerate the pace of molecular evolution (Fig.1); a larger value of *B* indicates a strong negative effect on the rate of molecular evolution, and *vice versa*. One can further verify that Eq.(14) is reduced to Eq.(2) in the case of *B*=0 (no macro-perturbation barrier).

**Fig 1.**
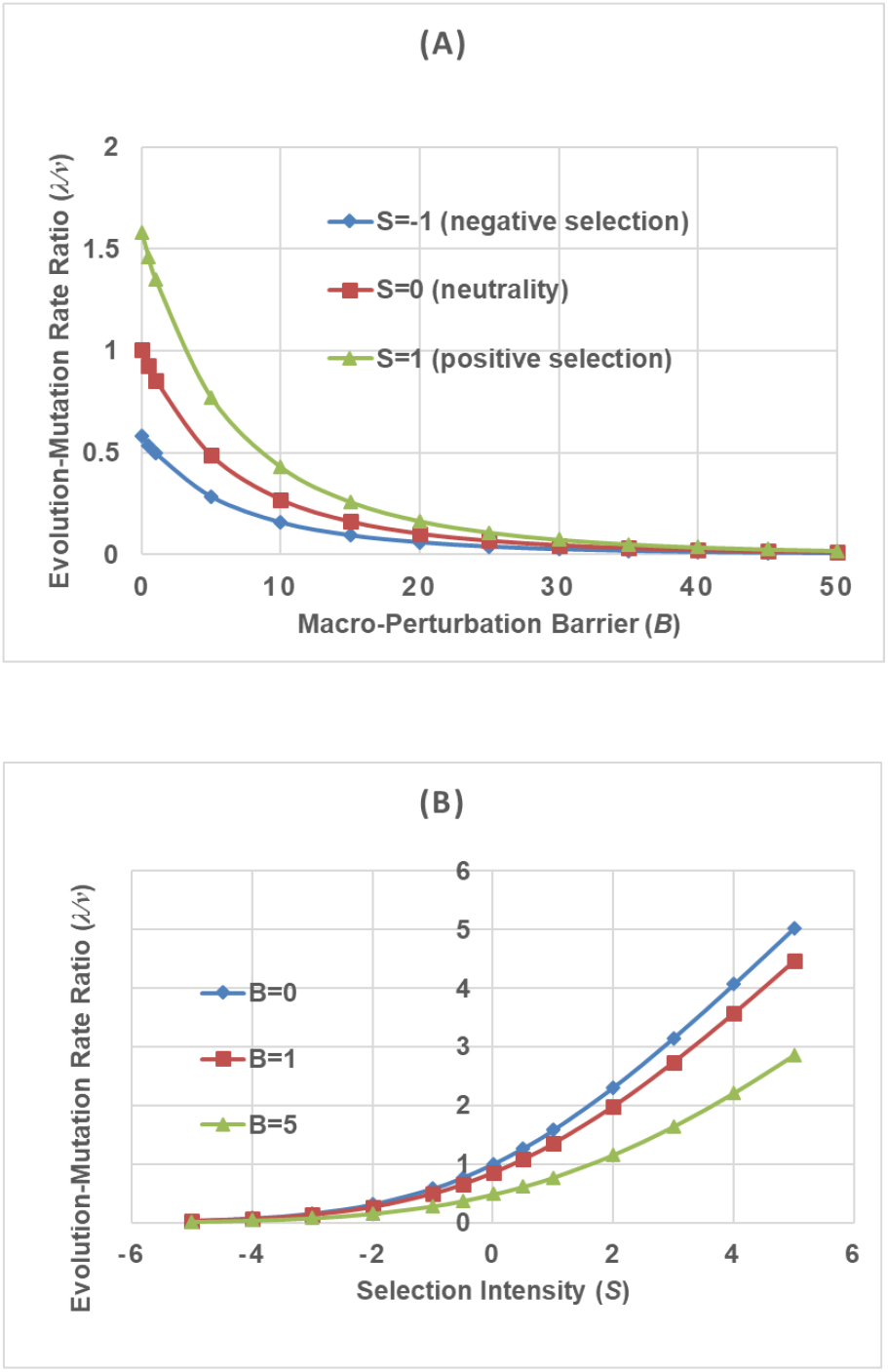
(A) The ratio of substitution rate to mutation rate (*λ/v*) plotted against the macro-perturbation barrier (*B*), given the selection intensity *S*=-1, *S*=0, or *S*=1, respectively. (B) The ratio plotted against the selection intensity (*S*), given the macro-perturbation barrier *B*=0, *B*=1 or *B*=5, respectively.

In the special case of neutral evolution (*S*=0), the evolutionary rate depends on the mutation rate (*v*) and macro-perturbation barrier (*B*), as given by

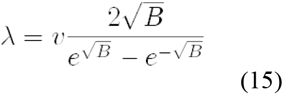

Apparently, Eq.(15) renders to the classical result of λ=*v* when *B*=0; otherwise, the rate of molecular evolution is always less than the mutation rate. On the other hand, when the magnitude of *S*, i.e., its absolute value, is large, those eigenvalues defined by Eq.(10) can be approximated by

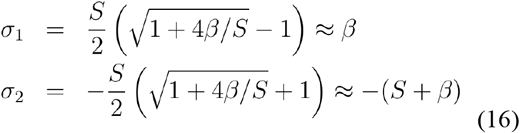

where *β* = *B*/*S*. Therefore, the rate of molecular evolution given by Eq.(14) can be approximated by

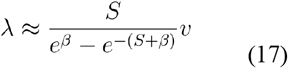

In the case of S ≫ 1, when the mutation is subject to a strong positive selection, Eq.(17) can be further simplified as *λ*/*v* ≈ *Se*^−*β*^. Hence, one may conclude that the rate of molecular evolution of a beneficial mutation can be reduced by the macro-perturbation barrier with a fact of *e*^−*β*^ that, roughly, depends on the perturbation-selection ratio *β* = *B*/*S* = *b*/(2*s*), but independent of the effective population size.

### Near-neutrality induced by macro-perturbation barrier

Eq.(15) claims that the evolutionary rate of a selectively-neutral mutant may inversely depend on the squared root of effective population size, 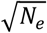, because of the macro-perturbation barrier. An interesting question is whether the pattern of nearly-neutral evolution emerges due to the macro-perturbation barrier. Consider the conventional criterion of near-neutrality, *N*_*e*_*s*∼ − 1. According to Eq.(2), the evolutionary rate-mutation rate ratio is *λ*/*v*∼0.0746. Equivalent to this criterion, by Eq.(15) we obtain B∼23.72. It means that, as long as the macro-perturbation barrier *B* is less than this value, a strictly-neutral mutant evolves like a nearly-neutral one.

### Quasi-neutrality under macro-perturbation barrier

Under the macro-perturbation barrier, the equivalence between the generic neutrality (*s*=0) and the golden standard of neutral evolution (*λ=v*) is no longer valid. In other words, to reach the golden standard of neutral evolution, the mutant must be beneficial (*s*>0). Hence, it may be appropriate to call the condition of *λ=v* the *quasi-neutrality*, as the result of the balance between the positive selection of the mutation and the macro-perturbation barrier. Letting *λ=v* in Eq.(14), we obtain

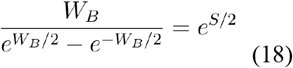

Eq.(18) shows that the condition of quasi-neutrality (λ=*v*) depends on the relative magnitude between *S* and *B*, but the *S-B* relationship is not straightforward. Intriguingly, after careful numerical analysis, we found that, for a broad range of *S* and *B*, Eq.(18) can be approximated by the following simple relationship

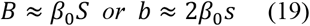

where *β*_0_ ≈ 3.148 (Fig.2). That is, the condition of quasi-neutrality is almost independent of the effective population size.

**Fig 2.**
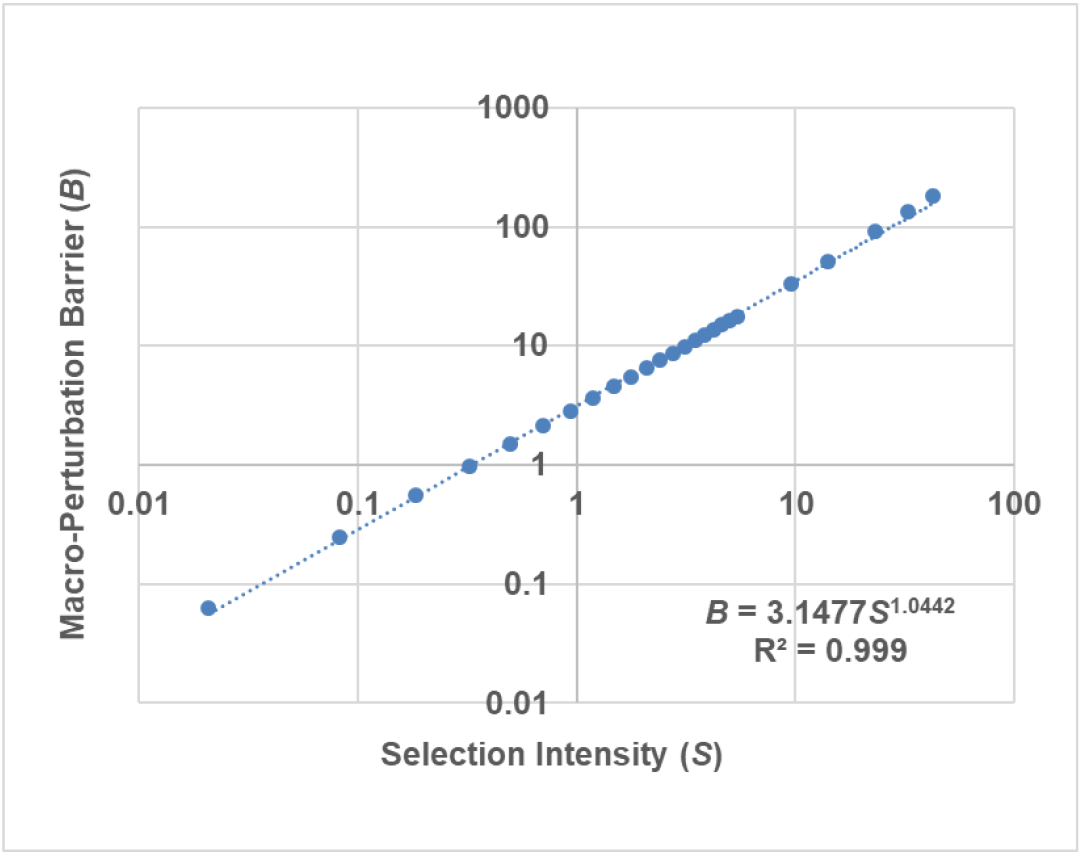
The relationship between the macro-perturbation barrier (*B*) and the selection intensity under the condition of quasi-neutrality (λ=*v*). This *B-S* relationship can be approximated by a power relationship, which can be further simplified as a linear relationship because the power component is very close to one.

### Selection-duality between generic neutrality and quasi-neutrality

In the case of generic neutrality (*s*=0), the rate of molecular evolution (*λ*) is given by Eq.(15), which is less than the mutation rate (*v*). Hence, a generically neutral mutation is subject to a negative selection because of the macro-perturbation. On the other hand, a mutant is under quasi-neutrality (*λ=v*) when *s* ≈ 0.159*b* according to Eq.(19), which means that this mutant must be slightly beneficial. These facts imply that quasi-neutrality actually requires at least slightly beneficial mutations, challenging the traditional wisdom of neutral theory. Intriguingly, a novel phenomenon called *selection-duality*, i.e., 0 < *s* < 0.159B, emerges between the generic neutrality and the quasi-neutrality: a slightly beneficial mutation defined above is subject to a negative selection, resulting in *λ*<*v*. The arising of selection-duality is apparently due to the negative effect of macro-perturbation on the fixation probability. In short, the generic neutrality and the fixation neutrality are the two boundaries for selection-duality.

### An evolutionary scenario of macro-perturbation

The mean fixation time *T*_*fix*_ is the number of generations, on average, required for a mutant to be fixed in a population (Crow and Kimura 1970; Ewens 2004). Meanwhile, the waiting time *T*_*wait*_ is the number of generations until such a mutation arises. The basic scenario of molecular evolution depicted by Kimura (1983) is valid only when this mean fixation time *T*_*fix*_ is much less than the waiting time *T*_*wait*_ before such a mutation arises. Consequently, molecular evolution between species can be unambiguously separated from the stage of population genetics, which can be modeled by a Markov-chain process. Note that the waiting time *T*_*wait*_ is technically determined by the inverse of the rate of molecular evolution. It appears that macro-perturbation increases the waiting time. For instance the waiting time of strict neutrality (no macro-perturbation) is 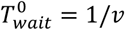 (*v* is the mutation rate), by Eq.(15) we have

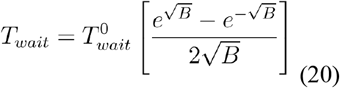

The mean fixation time (*T*_*fix*_) under macro-perturbation can be solved by the KAC backward equation but the analytical result is not available. Nevertheless, some preliminary numerical analyses showed that *T*_*fix*_ was affected by the macro-perturbation only marginally. Though a depth of analysis is needed, it is intuitively understandable as *T*_*fix*_ is the mean time that the mutant frequency reaches 1 before 0, conditional of no killing occurred.

### Fixation process of polyploidization: an illustration

Though the goal of the current study focuses on the theoretical formulation and perspective, it is desirable to illustrate how the theory of macro-perturbation can be useful to study the real evolutionary scenario. Here we discuss how this theory describe the fixation process of polyploidization: a balance between the short-term selective advantage and the genome instability.

Polyploidy, or the duplication of entire genomes (WGD), has been observed in prokaryotic and eukaryotic organisms. The consequences of polyploidization are complex, variable and often different greatly between species. However, the paucity of ancient polyploidy events that have ‘survived’ during the long term evolution suggested that polyploidy might be usually an evolutionary ‘dead end’ (Arrigo and Barker 2012; Vanneste et al. 2014; Mayrose et al. 2014). This is because polyploidy can have detrimental effects on fertility and fitness owing to genomic instability, mitotic and meiotic abnormalities, and gene expression and epigenetic changes, as been extensively discussed by and reviewed elsewhere (Comai, 2005).

On the other hand, there seems to be accumulating evidence that links short-term polyploid establishment and survival to periods of environmental and ecological upheaval. The notion of adaptive potential of polyploids attempted to explain why some new polyploids might be able to survive and even thrive in the short term (Soltis et al. 2009; te Beest et al. 2012; Madlung, 2013; Leitch and Leitch, 2008; Bomblies and Madlung 2014). Van de Peer et al. (2017) gave a comprehensive review for the underlying mechanisms, concluding that most explanations of the short-term success of polyploids are centered on the effects of genomic changes and increased genetic variation which are mediated by changes in gene expression and epigenetic remodelling (Lavania et al. 2012; Shi et al. 2012; Song and Chen 2015; Soltis et al. 2014; Parisod et al. 2010; Zhang et al. et al. 2016; Doyle et al. (2008); Schoenfelder and Fox 2015). Increased genetic variation can potentially affect the morphology, physiology and ecology of newly formed polyploids (te Beest et al. 2012; Soltis et al. 2014; McCarthy et al. 2016). Moreover, increased genetic variation in polyploids might also lead to increased tolerance to a broader range of ecological and environmental conditions, e.g., it has repeatedly been proposed that polyploids have a higher stress tolerance (environmental robustness) than do diploids, although this theory remains controversial (Mable et al.2011; Van de Peer et al. 2009; te Beest et al. 2012; Hahn et al. 2012).

Van de Peer et al. (2017) concluded that polyploidy is usually an evolutionary dead end, except in special circumstances when polyploids might have some selectively advantages over non-polyploids. According to the theory of macro-perturbation barrier of molecular evolution, polyploids usually carry a large macro-perturbation barrier. Consequently, frequency increase of polyploids in a population would be hindered randomly in each generation. In other words, the fixation probability of polyploids would be reduced considerably by the macro-perturbation barrier. Tentatively, one may use the quasi-neutrality as a justification for the fate of a polyploidy: the evolutionary rate of polyploids is greater than the arising rate of polylpoids when the selection coefficient of polyploids (s) exceeds a threshold given by Eq.(19), i.e., *s* > 0.159*b*; in this case polyploids has been strongly selected, ultimately toward the fixation of the population. While this criterion is independent of the effective population size, the larger is the population, the more efficient is the selection. When 0 < *s* < 0.159*b*, i.e., the case when polyploids are weakly adaptive, the evolutionary rate of polyploids is less than the arising rate of polylpoids. In short, the macro-perturbation barrier theory provides a population genetic foundation that helps to explain the apparently contradictory evolutionary fates of polyploids (Van de Peer et al. 2017).

### Concluding remarks

Molecular evolution describes how a new mutation can spread in a population, ultimately toward being fixed rather than being lost, due to either selection or genetic drift. Meanwhile, macro-perturbation refers to a rare, randomly occurred major genetic, epigenetic or environmental event that virtually stopped the fixation process. In this article, this phenomenon was modeled by a diffusion process with a killing (also called KAC process). We focused on the stochastic trajectory toward fixation of a mutation in a given population and showed that macro-perturbation can considerably reduce the rate of molecular evolution across species; for instance, a strictly-neutral mutant evolves as if a nearly-neutral under macro-perturbation. Moreover, we found a simple condition for the emergence of ‘quasi-neutrality’, i.e., the rate of molecular evolution equals to the mutation rate.

It should be noticed that killing function can be designed for a given KAC process, even under the same diffusion model. For instance, a complement macro-perturbation can be modeled by the KAC loss process of a mutant, that is, a loss process of this mutant could be killed at random time *τ* with a certain probability. Consequently, macro C-perturbation could indirectly increase the fixation probability, and so as the rate of molecular evolution. A more sophisticated scenario is that the rate of molecular evolution of a mutant could be affected by macro perturbation and C-perturbation simultaneously. We will address these more advanced issues in our future study.

